# Ornithological and molecular evidence of a reproducing *Hyalomma rufipes* population under continental climate in Europe

**DOI:** 10.1101/2023.01.13.523891

**Authors:** Gergő Keve, Tibor Csörgő, Anikó Benke, Attila Huber, Attila Mórocz, Ákos Németh, Béla Kalocsa, Enikő Anna Tamás, József Gyurácz, Orsolya Kiss, Dávid Kováts, Attila D. Sándor, Zsolt Karcza, Sándor Hornok

## Abstract

**Background:** Reports on adult *Hyalomma* ticks in certain regions of the Carpathian Basin date back to the 19^th^ century. These ticks were thought to emerge from nymphs dropping from birds, then molting to adults. Although the role of migratory birds in carrying ticks of this genus is known from all parts of Europe, in most countries no contemporaneous multiregional surveillance of bird-associated ticks was reported which could allow the recognition of hotspots in this context.

**Methods:** Ixodid ticks were collected from birds at seven ringing stations in Hungary, including both the spring and autumn migration period in 2022. *Ixodes* and *Haemaphysalis* species were identified morphologically, whereas *Hyalomma* species molecularly.

**Results:** From 38 passeriform bird species 957 ixodid ticks were collected. The majority of developmental stages were nymphs (n=588), but 353 larvae and 16 females were also present. On most birds (n=381) only a single tick was found and the maximum number of ticks removed from the same bird was 30. Tick species were identified as *Ixodes ricinus* (n=598), *Ixodes frontalis* (n=18), *Ixodes lividus* (n=6), *Haemaphysalis concinna* (n=322), and *D. reticulatus* (n=1). All twelve *Hyalomma* sp. ticks (11 engorged nymphs and an unengorged larva) were identified as *Hyalomma rufipes* based on three mitochondrial markers. This species was only found in the Transdanubian region and along its southeastern border. The Common Blackbird (*Turdus merula*) and the European Robin (*Erithacus rubecula*) were the two main hosts of *I. ricinus* and *I. frontalis*, whereas *H. concinna* was almost exclusively collected form long-distance migrants. The predominant hosts of *H. rufipes* were reed-associated bird species, the Sedge Warbler (*Acrocephalus schoenobaenus*) and the Bearded Reedling (*Panurus biarmicus*), both harboring these ticks at the end of June (i.e., the nesting period) in southwestern Hungary.

**Conclusions:** This study provides ornithological explanation for the regional, century-long presence of adult *Hyalomma* ticks under continental climate in the Transdanubian Region of the Carpathian Basin. More importantly, the autochthonous occurrence of a *H. rufipes* population was revealed for the first time in Europe, based on the following observations: (1) the bird species infested with *H. rufipes* are not known to migrate during their nesting period; (2) one larva was not yet engorged; (3) the larva and the nymphs must have belonged to different local generations; and (4) all *H. rufipes* found in the relevant location were identical in their haplotypes based on three maternally inherited mitochondrial markers, probably reflecting founder effect. This study also demonstrated that the species of ticks carried by birds were significantly different between collection sites even within a geographically short distance (200 km). Therefore, within a country multiregional monitoring is inevitable to assess the overall epidemiological significance of migratory birds in importing exotic ticks, and also in maintaining newly established tick species.

## Background

In the temperate zone of Europe, pathogens transmitted by hard ticks (Acari: Ixodidae) are responsible for the majority of the vector-borne diseases [1]. On this continent approx. 55 ixodid species occur [2]. From among these, the number of tick species that are regarded as indigenous will likely increase in several countries, in part due to climate change and the emergence of new, thermophilic tick species from the south.

In this scenario, the first prerequisite for the establishment of new tick species in any region is their repeated introduction, for which a very important natural route is via bird migration. Migratory birds are long-known carriers of ticks, most importantly *Hyalomma* species, from the south to temperate regions of Europe [3], even its northernmost parts [4]. However, birds usually carry immature ticks, larvae and nymphs of *Hyalomma* species [5], therefore in case of these thermophilic ticks, another crucial prerequisite prior to establishment is the ability of nymphs detaching from birds to molt to adults. This was already reported for both *Hyalomma marginatum* and *Hyalomma rufipes* from several countries north of the Mediterranean Basin, as exemplified by the UK [6] and the Netherlands [7] in western Europe, or Hungary in central Europe [8]. Consequently, *Hyalomma* adults might also overwinter [9], increasing the chances for future establishment of permanent, reproductive populations.

Recently, the emergence of *Hyalomma marginatum* was reported in a previously non-endemic region of the Mediterranean Basin in southern France, but it was stated that even in such newly invaded areas this tick species probably remains exclusively Mediterranean and cannot expand outside this climatic range [10]. On the other hand, north of the Mediterranean region, in the Carpathian Basin (geographically including both Hungary and the Transylvanian Basin: [11]), adult ticks from the genus *Hyalomma* are long-known for their autochthonous occurrence under continental climate. This was already reported in the 19th century [12], and later confirmed [13,14]. At the same time, in the absence of detailed morphological description, the species in the Carpathian Basin remained uncertain, because some hints were more relevant to *H. rufipes* (e.g., the name *Hyalomma aegyptium*: [14]), while others to *H. marginatum* (as implied in the predominance of the species referred to from Hungary in the Mediterranean Basin: [13]). More recently, *H. rufipes* adults were found on cattle on two occasions in Hungary [8], and one adult on the same host species 10 years later by citizen science method [15].

Interestingly, these century-long reports on the presence of adult *Hyalomma* ticks in the Carpathian Basin attest that the chance for their occurrence is more likely in certain endemic areas of the country. However, this hypothesis was not yet tested from the point of view of bird migration, despite the long-known import of *Hyalomma* nymphs by birds into this geographical region [16]. In light of the above, the aim of this study was to perform a pilot survey focusing on the comparison of tick species carried or imported by birds at various locations in Hungary. These locations were meant to represent most of the Carpathian Basin where important stopover sites can be found along the Adriatic Flyway of bird migration.

## Methods

### Sample collection

In this study, birds mist-netted at seven ringing stations (Figure 1) by standard ornithological mist-nets (mesh size 16 mm) were examined for the presence of ticks, between March and November, 2022. The main characteristics of ringing stations are as follows:

**Figure 1.**
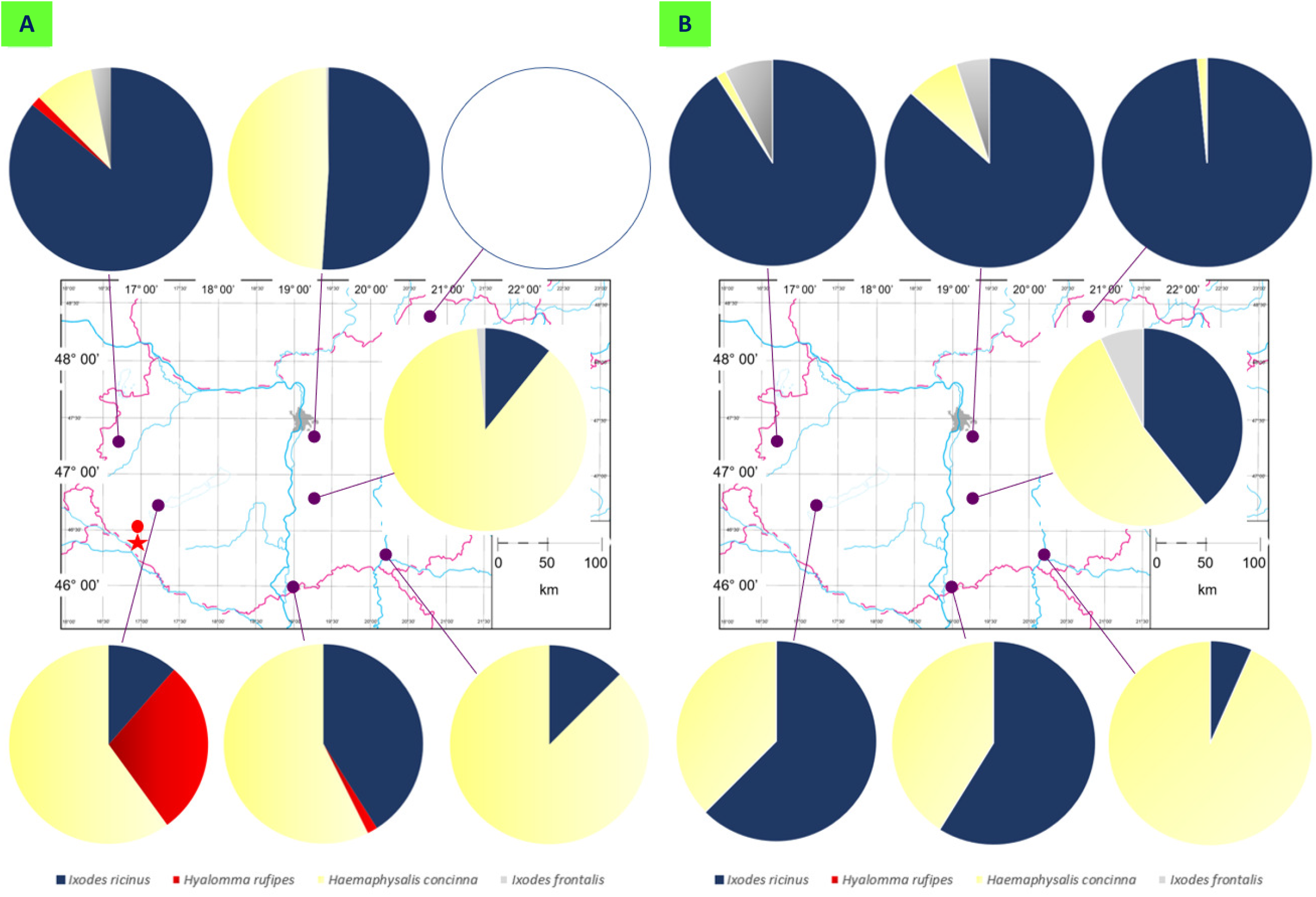
Map of Hungary showing ringing stations and the ratio of tick species collected in (A) the first semiannual period (March to July) and (B) the second semiannual period (August to November). In the former (A) the location of the first *Hyalomma* nymph reported from a bird in Hungary in 1955 is marked with a red dot, and the place where adult *Hyalomma rufipes* ticks were found on cattle is indicated with a red star.

1. Tömörd Bird Ringing Station (coordinates: 47°21’N, 16°39’E): situated in northwestern Hungary, next to a small lake. It is surrounded by cultivated lands, bushes and deciduous forest, predominantly oak trees.
2. Ócsa Bird Ringing Station (47°19’N, 19°13’E): situated in north central Hungary, on the edge of a wetland. It is surrounded by arable fields, poplar plantations with several interspersed open-pit gravel mines [17].
3. Bódva Valley Bird Ringing Station (coordinates: 48°27’N, 20°42’E): situated in northeastern Hungary, and located in the valley of Bódva River. The river is surrounded by the mosaics of gallery forests, wet meadows, *Prunus* scrubs and arable lands. The adjacent hillsides are covered mostly by oak forests.
4. Fenékpuszta Bird Ringing Station (46°44’N, 17°14’E): situated in southwestern Hungary, next to the largest lake in Central Europe, Lake Balaton. The vegetation type is predominantly reed.
5. Izsák, Lake Kolon Bird Ringing Station (coordinates: 46°46’N, 19°19’E): situated in central Hungary, across Lake Kolon in the reedbed. The vegetation type is typical for marshes, heavily covered with reeds. It is surrounded by cultivated lands and planted forests.
6. Dávod, Lake Földvár Bird Ringing Station (coordinates: 46°0’N, 18°51’E): situated in south central Hungary on the shores of Lake Földvár which is an oxbow lake, that was formed naturally from river Danube. The lake is surrounded by reedbed. Sand Martins (*Riparia riparia*) were ringed in the sand mines of Baja (coordinates: 46°12N, 18°58’E) which are close to this area.
7. Lake Fehér Ornithology Camp (coordinates: 46°20’N, 20°6’E). The camp is located near to Lake Fehér, which is large fishpond system the greatest saline lake. The vegetation type is reedbed with sparse shrubs and trees.

Ticks were removed from the skin of birds with fine tweezers and stored in 96% ethanol. Data of collection (date, location, avian host species, ring number) were recorded. *Ixodes* and *Haemaphysalis* species were identified morphologically [2], whereas *Hyalomma* species molecularly as outlined below.

For data comparison and presentation, ornithological traits were assigned to bird species according to Csörgő et al. [18]. English bird species names are capitalized in accordance with the international recommendations (https://bou.org.uk/british-list/bird-names/).

### DNA extraction

Ticks of the genus *Hyalomma* were disinfected on their surface with sequential washing for 15 s in 10% NaClO, tap water and distilled water. For the DNA extraction, the larva was used without incision, whereas nymphs were cut dorsally on the idiosoma. DNA was extracted with the QIAamp DNA Mini Kit (QIAGEN, Hilden, Germany) according to the manufacturer’s instruction, including an overnight digestion in tissue lysis buffer and Proteinase-K at 56 °C. Extraction controls (tissue lysis buffer) were also processed with the tick samples to monitor cross-contamination.

### Molecular identification of *Hyalomma* species

The cytochrome oxidase subunit I (*cox*1) gene was chosen as the first target for molecular analysis. The PCR was modified from Folmer et al. [19] and amplifies an approx. 710-bp-long fragment of the gene. The primers HCO2198 (5’-TAA ACT TCA GGG TGA CCA AAA AAT CA-3’) and LCO1490 (5’-GGT CAA CAA ATC ATA AAG ATA TTG G-3’) were used in a reaction volume of 25 μl, containing 1 U (stock 5 U/μl) HotStarTaq Plus DNA Polymerase, 2.5 μl 10× CoralLoad Reaction buffer (including 15 mM MgCl_2_), 0.5 μl PCR nucleotide Mix (stock 10 mM), 0.5 μl of each primer (stock 50 μM), 15.8 μl ddH2O and 5 μl template DNA. For amplification, an initial denaturation step at 95 °C for 5 min was followed by 40 cycles of denaturation at 94 C for 40 s, annealing at 48 °C for 1 min and extension at 72 °C for 1 min. Final extension was performed at 72 °C for 10 min.

Another PCR was used to amplify an approx. 460-bp-fragment of the 16S rDNA gene of Ixodidae [20], with the primers 16S+1 (5’-CTG CTC AAT GAT TTT TTA AAT TGC TGT GG-3’) and 16S-1 (5’-CCG GTC TGA ACT CAG ATC AAG T-3’). Other reaction components, as well as cycling conditions were the same as above, except for annealing at 51 °C. In addition, a conventional PCR reaction was used with the primer pairs T1B (5’-AAA CTA GGA TTA GAT ACC CT-3’) and T2A (5’-AAT GAG AGC GAC GGG CGA TGT-3’) to amplify an approx. 360-bp-long fragment from the 12S rRNA gene from all DNA extracts [21,22]. The PCR was modified with the following conditions. An initial denaturation step at 95 °C for 5 min was followed by 5 cycles of denaturation at 94 °C for 30 s, annealing at 50 °C for 30 s and extension at 72 °C for 30 s and 30 cycles of denaturation at 94 °C for 30 s, annealing at 53 °C for 30 s and extension at 72 °C for 30 s. Final extension was performed at 72 °C for 7 min [22,23].

### Sequencing

In all PCRs non-template reaction mixture served as negative control. Extraction controls and negative controls remained PCR negative in all tests. Purification and sequencing of the PCR products were done by Biomi Ltd. (Gödöllő, Hungary). Quality control and trimming of sequences were performed with the BioEdit program, then alignment with GenBank sequences by the nucleotide BLASTN program (https://blast.ncbi.nlm.nih.gov). New sequences were submitted to GenBank (*cox*1: OQ108291-OQ108294, 16S rRNA: OQ103402-OQ103405, 12S rRNA: OQ103398-OQ103401).

### Statistical analyses

Fisher exact test was used to compare prevalence rates and differences were regarded significant if P < 0.05.

## Results

### (1) Species and developmental stages of ticks infesting birds

During 2022, 540 individuals of 38 passeriform bird species were found to be tick-infested, from which altogether 957 ixodid ticks were collected. The majority of developmental stages were nymphs (n=588), but 353 larvae and 16 females were also present. On most birds (n=381) only a single tick was found. The maximum number of ticks removed from a single bird was 30, and the mean intensity of tick-infestation was 1.78 tick/tick-infested bird in the whole study period.

Based on morphological characteristics, the ticks belonged to the following species: *Ixodes ricinus* (n=598), *Ixodes frontalis* (n=18), *Ixodes lividus* (n=6), *Haemaphysalis concinna* (n=322), and *D. reticulatus* (n=1) (Supplementary Table 1). Morphologically, the twelve *Hyalomma* sp. ticks could only be identified on the genus level and their molecular identification was necessary. All *Hyalomma* nymphs were in a similar, advanced state of engorgement, but the single larva was flattened, apparently unengorged.

Based on the 16S rRNA gene, *Hyalomma* nymphs belonged to three haplotypes (Table 1). One of these collected in south Hungary (OQ103402) had 100% (383/383 bp) sequence identity to *H. rufipes* previously collected from a bird in north-central Hungary (Ócsa: KU170517) and another in Egypt (MK737650). The second haplotype (collected in northwest Hungary: OQ103403) differed in two, and the third haplotype (all other specimens: OQ103404-OQ103405) in one position of their 16S rRNA sequence, meaning 99.5% and 99.7% sequence identities to the above two reference sequences, respectively (Table 1). One haplotype (OQ108291) differed in one position, whereas all other *H. rufipes* specimens were 100% (645/645 bp) identical in the sequenced part of their *cox*1 gene (OQ108292-OQ108294) to a tick collected from Eurasian Reed Warbler (*Acrocephalus scirpaceus*) in the Netherlands (MT757612) and another reported from Malta (OL339477). Interestingly, these *cox*1 sequences were even more different (in two bps) from *H. rufipes* collected from a bird in a previous study in north-central Hungary (Ócsa: KU170491). In addition, all *H. rufipes* nymphs and the larva had identical 12S rRNA sequences (OQ103398-OQ103401), with 100% (341/341 bp) sequence identity to ticks collected from birds in Malta (OL352890) and in Italy (MW175439). Thus, the genus *Hyalomma* was exclusively represented by *H. rufipes* (n=12).

**Table 1.** Data of Hyalomma ticks collected in 2022 from birds at various ringing stations in Hungary. Identical background color in cells of the same column of a genetic marker indicates identical sequences.

| Isolate code | Bird species | Date | Region of Hungary (location) | *Hyalomma* sp. (number, stage) | GenBank accession numbers according to the three genetic markers |  |  |
| --- | --- | --- | --- | --- | --- | --- | --- |
|  |  |  |  |  | 16S rRNA | Cox1 | 12S rRNA |
| BA2 | SYL COM | May 14 | south (Dávod) | *H. rufipes* (1×N) | OQ103402 | OQ108291 | OQ103398 |
| GJ10 | FIC HYP | April 23 | northwest (Tömörd) | *H. rufipes* (1×N) | OQ103403 | OQ108292 | OQ103399 |
| BE02 | ACR SCH | June 26 | southwest (Fenékpuszta) | *H. rufipes* (1×N) | OQ103404 | OQ108293 | OQ103400 |
| BE03 | ACR SCH | June 26 | southwest (Fenékpuszta) | *H. rufipes* (1×N) | OQ103404 | OQ108293 | OQ103400 |
| BE04 | ACR SCH | June 26 | southwest (Fenékpuszta) | *H. rufipes* (1×N) | OQ103404 | OQ108293 | OQ103400 |
| BE05 | PAN BIA | June 26 | southwest (Fenékpuszta) | *H. rufipes* (1×N) | OQ103405 | OQ108294 | OQ103401 |
| BE06 | PAN BIA | June 26 | southwest (Fenékpuszta) | *H. rufipes* (1×L, 5×N) | OQ103405 | OQ108294 | OQ103401 |

**Abbreviations:**
ACR SCH = *Acrocephalus schoenobaenus*, FIC HYP = *Ficedula hypoleuca*, PAN BIA = *Panurus biarmicus*, SYL COM = *Sylvia communis*

### (2) Host-associations of tick species and the migratory habits of their avian hosts

Associations of ticks collected in this study with different bird species are summarized in Supplementary Table 2. The Common Blackbird (*Turdus merula*) (n=58) and the European Robin (*Erithacus rubecula*) (n=105) were the two main hosts of *I. ricinus* in both the spring and the autumn tick collection periods. The preferred hosts of *I. frontalis* were also these two bird species (n= 4 and 5, respectively). *Haemaphysalis concinna* most often infested the Sedge Warbler (*Acrocephalus schoenobaenus*) (n=49) and Savi’s Warbler (*Locustella luscinioides*) (n=65) (Supplementary Table 2). *Hyalomma rufipes* was only collected on repeated occasions from Sedge Warblers (*A. schoenobaenus*) and Bearded Reedling (*Panurus biarmicus*), and once from a Common Whitethroat (*Curruca communis*) and from a European Pied Flycatcher (*Ficedula hypoleuca*). *Ixodes lividus* was only found once, on its specific host, the Sand Martin (*Riparia riparia*). Importantly, with the exception of the accidental finding of a single *D. reticulatus* female on a Common Blackbird, all other females (n=16) belonged to the two ornithophilic tick species *I. frontalis* (n=9) and *I. lividus* (n=6).

During the spring, at the ringing station in north-central Hungary (Ócsa) where the highest number of tick-infested birds were caught and which contributed the most balanced ratio of birds with different migratory habits to the study, there was a highly (P < 0.0001) significant difference between the host associations of *I. ricinus* and *H. concinna*, since the former predominated on resident and short-distance migrant bird species, but *H. concinna* on long-distance migrants. In the autumn, taking into account all ringing stations, the difference between these two tick species in the same comparison, and the association of *I. frontalis* with resident and short-distance migrant bird species was also highly significant (P < 0.0001).

### (3) Spatiotemporal occurrence of tick species

*Ixodes ricinus* and *H. concinna* were found to infest birds in both the spring and autumn collection periods (Figure 1), whereas the presence of *H. rufipes* was restricted to the first half of the year (Figure 1, Table 1), and *I. frontalis* predominated in the autumn period (Figure 1). Importantly, *H. rufipes* was collected from long-distance migrant birds in south and northwest Hungary in May and April, respectively (Table 1). However, all other specimens of this species were removed from birds in the middle of summer (late June) at one ringing station in the southwestern part of the country (Fenékpuszta).

During the spring period, *I. ricinus* was the predominant tick species in the north, whereas *H. concinna* in central and south Hungary (Figure 1, Supplementary Table 1). However, in the autumn, *I. ricinus* represented the highest number of ticks from birds in north as well as in southwestern parts of the country, and *H. concinna* at two ringing stations, in central and southeast Hungary (Figure 1). *Hyalomma rufipes* was only found in the Transdanubian region and in one case along the southern reach of the Danube. On the other hand, *I. frontalis* could only be collected in northern and central locations during both spring and autumn and was absent from birds in southern parts of the country (Figure 1).

Taken together, *I. ricinus* and *H. concinna* occurred on birds at all sampling sites, but their ratio was different according to these sites and semiannual periods. At the same time, the spatiotemporal distribution was limited in case of *H. rufipes* and *I. frontalis*.

## Discussion

In Hungary, studies on tick-infestations of birds date back to more than half a century [16], and have been ever since extensively performed on annual or tri-annual bases focusing on the same ringing station in the north-central part of the country (Ócsa: [24–26]). Similar reports on ticks from avian hosts are available from numerous European countries, as exemplified by Sweden [27], The Netherlands and Belgium [28], Germany [29] or Italy [30]. Relevant studies have also been reviewed recently [31,32]. However, discounting opportunistic and sporadic collections of ticks from birds, the present study is the first “horizontal tick survey” from birds in the Carpathian Basin and probably also in a broader geographical context. This implies that ticks were removed and their species identified at several ringing stations simultaneously in the course of one year, allowing not only the regional comparison of tick burdens carried by birds, but also assessing the significance and need of similar studies on a larger, continental scale.

In this study, six species of ixodid ticks (three prostriate and three metastriate) were collected from birds. The most significant finding related to tick species diversity was the *H. rufipes*-infestation of three long-distance migrant and a resident bird species. Importantly, *H. rufipes* was collected in south and northwestern Hungary during late spring in 2022, as in a previous study [26]. However, in this study all remaining 10 specimens were removed from birds in the middle of summer (late June) at one ringing station in the southwestern part of the country (Fenékpuszta), i.e., in the same county (Zala) where *Hyalomma*-infestation of a bird was diagnosed for the first time in Hungary in 1955 [16]. In the same region, *Hyalomma* sp. ticks were reported to occur [33] and *H. rufipes* adults were identified on two occasions from cattle [8] (Figure 1.A).

It is utterly unlikely that all five individuals of the two avian host species of these 9 fully engorged nymphs and one unengorged larva of *H. rufipes* (sampled on June 26) carried these ticks into Hungary from abroad. *Hyalomma rufipes* has a two-host life cycle, and engorged nymphs drop off from the host after 21-29 days of infestation [34]. One of the avian hosts shown to harbor nymphs of *H. rufipes* in this study, the Sedge Warbler (*A. schoenobaenus*) typically arrives in Hungary from the wintering grounds in Africa between April and early May [18], and late June (when its *Hyalomma*-infestation was diagnosed) is in the middle of its nesting period, without migration. On the other hand, the other repetitive host of *H. rufipes* in this study, the Bearded Reedling (*P. biarmicus*) is an *a priori* resident bird species, with rarely documented limited movements, but according to ringing data [18] these “vagrancies” never occur in its summer nesting period.

Regarding the results of molecular analyses, it is not surprising that all *H. rufipes* individuals collected in 2022 from birds in Hungary (n=12) had identical 12S rRNA haplotypes, because this genetic marker was shown to be identical in case of a much larger set of *H. rufipes* ticks (n=48) collected from birds with probably different geographical origin [35]. However, in this study the sequenced part of the *cox*1 gene was also identical between all *H. rufipes* (n=11) collected in the Transdanubian part of Hungary, in particular in case of those 10 ticks which were removed from birds at the same ringing station in southwest Hungary (Fenékpuszta). *Hyalomma rufipes* was shown to differ remarkably in its *cox*1 haplotype in case of ticks carried by birds with different geographical origin [35]. Moreover, the ratio and presence or absence of certain *Hyalomma cox*1 haplotypes were demonstrated to be site- and population-specific, usually with multiple haplotypes even within the same population [36]. Therefore, finding of exclusively one *cox*1 haplotype among 10 *H. rufipes* ticks collected in one location (Fenékpuszta) raises the possibility that these ticks represent the same population. Their genetic similarity is probably a consequence of founder effect. Taken together, all three studied mitochondrial, maternally inherited genetic markers were identical only between *H. rufipes* individuals collected in the latter place, also supporting the common maternal aborigine of these ticks.

In addition, the apparently unengorged state of the *H. rufipes* larva on one of these birds also argues against the foreign origin of its tick-infestation. Note that in a previous study only molting (i.e., advanced stage) *H. marginatum* larvae were found on birds in Hungary, and all other stages were nymphs [25,26]. Importantly, hitherto molecularly verified *H. rufipes* larvae were only reported from birds in south European countries (reviewed by Keve et al. [32]), and typically only nymphs of this tick species arrive on birds in countries north of the Mediterranean Basin if these originate from Africa (Figure 2; [32]).

**Figure 2.**
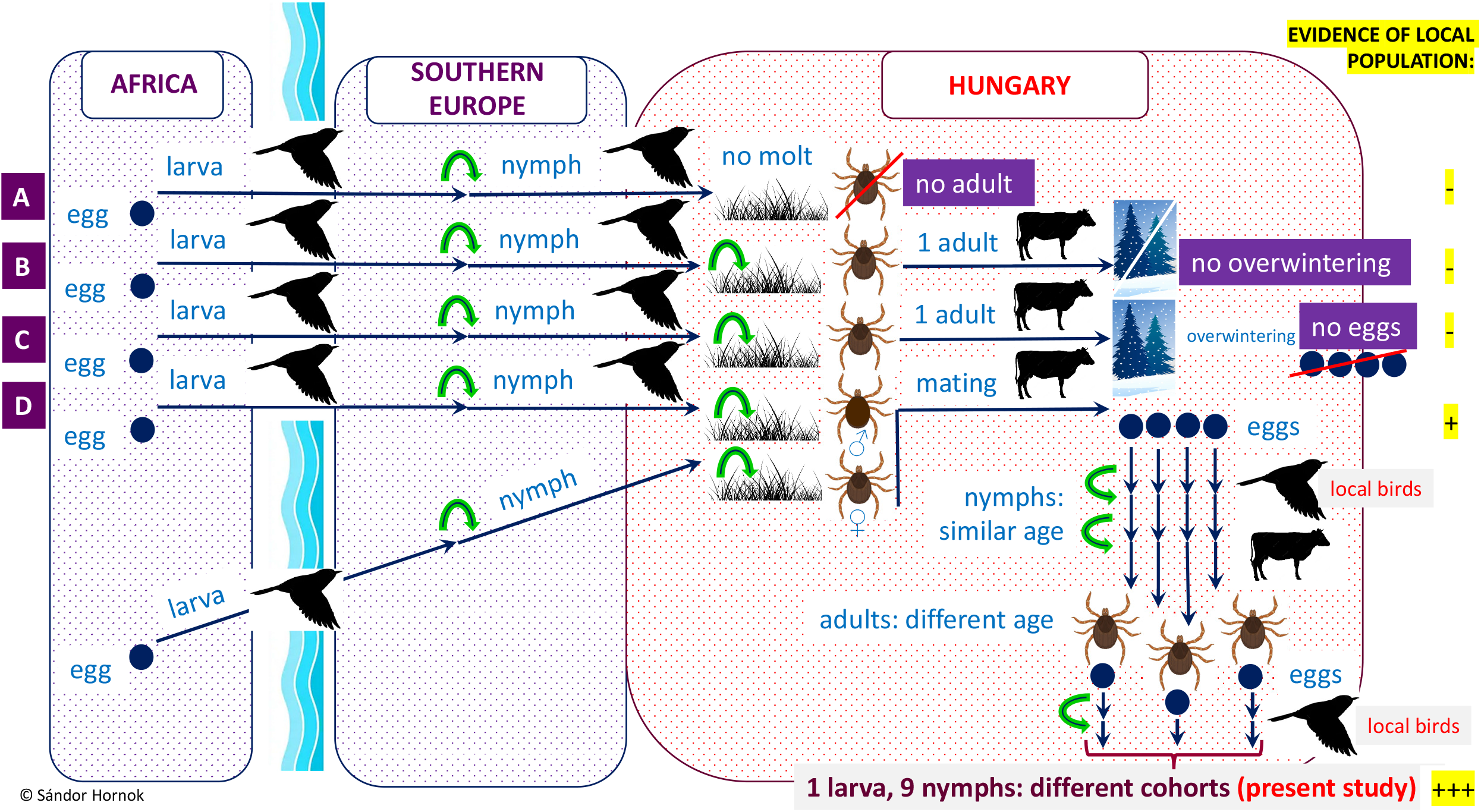
Illustration of the possible consequences of bird-borne transportation of *Hyalomma rufipes* into countries north of the Mediterranean Basin, including Hungary. Green arrows indicate molting. (A) Nymphs transported by birds may die after drop-off, or (B) molt to adult which cannot overwinter, or (C) if they overwinter as adults, females will not produce eggs in the absence of previous mating, or (D) if nymphs carried by birds detach and molt to male and another (carried independently) to female and these meet and mate on cattle, females will be able to lay eggs after drop-off. First generation larvae and nymphs developing from these eggs probably will have a similar state of engorgement but molting to adults they will find host and will mate at different time. Therefore, existence of a second generation may involve the simultaneous presence of larvae and nymphs of different cohorts on local birds, as shown in this study.

Obviously, not all ticks carried by migratory birds in the spring were imported by them from southern countries, and this is particularly relevant to those avian hosts which arrive from their wintering grounds during the activity peak of local tick populations. Similarly to previous bird tick studies in the Carpathian Basin [26] and most countries north of the Mediterranean Basin (e.g., [27]), *I. ricinus* was the tick species most commonly collected from birds in 2022 in Hungary. *Haemaphysalis concinna* was the second most abundant tick species on birds, which, however, seems to be unique to the Carpathian Basin and its region [32]. Both of these tick species (*I. ricinus, H. concinna*) indigenous to Hungary tend to infest birds which arrive in their main activity periods [37], therefore *I. ricinus* (peak activity: April) is mainly found on residents and short-distance migrants (typically arriving early spring), whereas *H. concinna* (peak activity: May) on long-distance migrants (usually arriving late spring) [18].

New tick-host associations revealed in this study include the presence of *H. rufipes* on the Bearded Reedling (*P. biarmicus*), and infestation of Moustached Warbler (*Acrocephalus melanopogon*) with *H. concinna*. Although *D. reticulatus* seldom occurs on birds [32], its immature developmental stages were reported from avian hosts (including the Common Blackbird, *T. merula*) [38]. The collection of its adult on a Blackbird during this study is probably an accidental finding.

Considering the regional occurrence of tick species on birds in the Carpathian Basin, *H. concinna* is a thermophilic tick species [39], and this is in accordance with its predominance on birds in central-south Hungary during the spring, and central-southeastern Hungary in the autumn (i.e., the warmest regions of the country: Supplementary Figure 1). On the other hand, the reason for the absence of *I. frontalis* from birds in the southern part of Hungary maybe twofold. First, the relevant sampling locations are near water surfaces where the predominant bird species (e.g., Savi’s Warbler, *L. luscinioides*) are not known to be hosts or (e.g., the Sedge Warbler, *A. schoenobaenus*) are exceptional hosts of this tick species [32]. Second, in these places bird mist-netting (i.e., tick collection) was terminated sooner than the late autumn peak activity of *I. frontalis* in the relevant region [40].

In this study, *H. rufipes* was only found in the Transdanubian region and once along the southern Danube, in line with the reported 130-year-long endemicity of *Hyalomma* species in the country [12–14]. While *Hyalomma*-infestation was previously reported on non-water-associated bird species (*E. rubecula, C. communis*) in the springtime in north-central Hungary (Ócsa) [25,26], this is the first occasion when ticks of this genus were observed on reed-dwelling birds in another region of Hungary, in a different season (during summer). This also raises the question on what the differences between the relevant two habitats in terms of landscape, vegetation and avian hosts are.

Fenékpuszta Bird Ringing Station is situated next to Lake Balaton. Here, the reedbed habitat in the riparian zone narrows to about 150 meters at the site of the mist-nets, where 12 pieces of these stretch across the reedbed completely. Due to uninterrupted reeds, this is an important stopover site for migrating passerines, particularly *Acrocephalus*-species. Based on ringing data, mostly long-distance migrant Sedge Warblers (*A. schoenobaenus*) and Eurasian Reed Warblers (*A. scirpaceus*) stop in this area, but Great Reed Warblers (*A. arundinaceus*) and Savi’s Warblers (*L. luscinioides*) are also significant in numbers.

Conversely, in Ócsa Bird Ringing Station the heterogeneous reedbed habitats of the capture locations are interspersed with fast growing shrubs as elderberry (*Sambucus nigra*) and blackberry (*Rubus fruticosus*), with softwood stands (*Salix* spp. and *Populus* spp.) form most of the vegetation. Thus, the Eurasian Blackcap (*Sylvia atricapilla*) and the European Robin (*Erithacus rubecula*) are two most common short-to-mid-distance migratory species here [17]. Regarding the capture rates of the species groups of migrating passerines, there is a significant difference between the homogeneous reedbed and other habitats (where the reedbed is patchy and alternates with deciduous forests, berry bushes). While *Acrocephalus* spp. account for the largest proportion of birds caught in Fenékpuszta, bush-dwelling warblers present a higher portion in Ócsa.

Based on the above, the existence of at least one indigenous population of *H. rufipes* is evidenced in the western part of Transdanubia, near Lake Balaton, because of the following reasons: (1) most importantly, the recognized avian hosts of *H. rufipes* were extremely unlikely to arrive from abroad shortly prior to their examination, especially not all five of them; (2) one larva was not yet engorged; (3) the larva and the nymphs (in a similar state of engorgement) were offspring of two females and must have belonged to different local generations (Figure 2); and (4) all *H. rufipes* found in the relevant location were identical in their haplotypes based on three maternally inherited mitochondrial markers, probably reflecting founder effect.

In addition, adults of *H. rufipes* are known to occur in the western part of the Carpathian Basin for 130 years, and in the same county (Zala) with its present collections adults of this tick species were found to infest cattle repeatedly [8]. Small local populations of *H. rufipes* were proposed to explain the occasional presence of *H. rufipes* in Russia [41,42] and its populations in scattered areas are also known in north Africa [42,43]. However, to our knowledge, this is the first report of a similar phenomenon and its evidence from Europe. The most important limiting factor for the survival of this xerophilic tick species under any climate is thought to be the maximum level of precipitation (annual rainfall) which is around 650 mm in southwestern Hungary (Supplementary Figure 1), i.e., similar to what is well-tolerated by *H. rufipes* in its range within Africa [44,45]. Populations of these ticks probably can survive winter conditions as adults in southwestern Hungary where winter temperatures are among the mildest in the country (Supplementary Figure 1). Nevertheless, *H. rufipes* is known to have populations in regions with up to 120 days of frost [42]. It is also noteworthy here that the likely overwintering of *H. rufipes* was reported in the Czech Republic [9], north of Hungary. Importantly, the discovered *H. rufipes* population might act as a “stepping-stone” for this tick species during its northward transportation by birds which use the relevant habitat near Lake Balaton in southwestern Hungary as a stopover site (see above).

On the other hand, no evidence was gained for any further *Hyalomma* populations indigenous in other regions of Hungary, as also indicated by the overall absence of *Hyalomma* ticks from birds in the autumn migration period. Thus, also taking into account the over-century-long presence of adult *Hyalomma* ticks, up to now there was no evidence for their emergence in the Carpathian Basin, but here evidence is reported for the emergence of a local population for the first time.

Similarly relevant to a broader, international context, the most important aim of the present study was also fulfilled, i.e., it was successful to demonstrate discrepancies between sampling sites, indicating that in the above context single-site surveys may be biased (not informative) on the actual risk posed by birds in transporting ticks in a geographical region or country. Therefore, to state the emergence or increasing presence of a *Hyalomma* species, ticks should be collected (larvae and nymphs from birds, and/or adults from reproductive hosts) extensively and annually in different regions of suspected endemic areas, preferentially by unbiased professionals who should stick to a standard methodology (sampling protocol).

## Abbreviation

*cox*1: cytochrome *c* oxidase subunit I

## Declarations

### Ethics approval

The study was carried out according to the national animal welfare regulations (28/1998). All songbirds were handled and released by experienced ringers of BirdLife Hungary.

License for bird ringing was issued by the Pest County Government Authority (/https://www.mme.hu/sites/default/files/pe_ktf_97_13_2017_vvt.pdf).

### Consent to participate

Not applicable.

### Consent for publication

Not applicable.

### Availability of data and materials

The sequences obtained during this study are deposited in GenBank under the following accession numbers: *cox*1 (OQ108291-OQ108294), 16S rRNA (OQ103402-OQ103405), 12S rRNA (OQ103398-OQ103401). All other relevant data are included in the manuscript and the references or are available upon request by the corresponding author.

### Authors’ contributions

GK: conceptualization, study design, sample collection, tick species identification, manuscript writing. TC, AB, AH, AM, ÁN, BK, EAT, JG, OK: study design, sample collection, data curation. DK: ornithological categorization, manuscript writing. ADS: study design, supervision. ZK: study organization, data availability. SH: conceptualization, study design, DNA extraction, molecular analyses, manuscript writing, preparation of figures.

## Acknowledgement

The authors are grateful to Ms. Nóra Takács for performing PCRs, to Dr. Jenő Kontschán for his advice in the terminology and definition of populations and to Németh Anna for her participation in field trips. The authors also thank all staff members and ringing personnel who contributed to the sample collection.

## Figure Legends

**Supplementary Figure 1.** The average annual precipitation (https://www.met.hu/) and temperature (https://www.mozaweb.com/search?search=középhőmérséklet) in January in Hungary, based on data from the Hungarian Meteorological Service (OMSZ). The site of the discovered *Hyalomma rufipes* population is marked with a star.

## References

1. Rochlin I, Toledo A. Emerging tick-borne pathogens of public health importance: a mini-review. J Med Microbiol. 2020;69:781–91.

2. Estrada-Peña A, Mihalca AD, Petney TN. Ticks of Europe and North Africa: a guide to species identification. Springer; 2017.

3. Hoogstraal H, Kaiser MN, Traylor MA, Gaber S, Guindy E. Ticks (Ixodoidea) on birds migrating from Africa to Europe and Asia. Bull World Health Organ. World Health Organization; 1961;24:197.

4. Brinck P, Svedmyr A, von Zeipel G. Migrating birds at Ottenby Sweden as carriers of ticks and possible transmitters of tick-borne encephalitis virus. Oikos. JSTOR; 1965;88–99.

5. Capek M, Literak I, Kocianova E, Sychra O, Najer T, Trnka A, et al. Ticks of the Hyalomma marginatum complex transported by migratory birds into Central Europe. Ticks Tick-Borne Dis. 2014;5:489–93.

6. Hansford KM, Carter D, Gillingham EL, Hernandez-Triana LM, Chamberlain J, Cull B, et al. Hyalomma rufipes on an untraveled horse: Is this the first evidence of Hyalomma nymphs successfully moulting in the United Kingdom? Ticks Tick-Borne Dis. Elsevier; 2019;10:704–8.

7. Uiterwijk M, Ibanez-Justicia A, van de Vossenberg B, Jacobs F, Overgaauw P, Nijsse R, et al. Imported Hyalomma ticks in the Netherlands 2018-2020. Parasit Vectors. London: Bmc; 2021;14:244.

8. Hornok S, Horváth G. First report of adult Hyalomma marginatum rufipes (vector of Crimean-Congo haemorrhagic fever virus) on cattle under a continental climate in Hungary. Parasit Vectors. 2012;5:170.

9. Rudolf I, Kejikova R, Vojtisek J, Mendel J, Penazziova K, Hubalek Z, et al. Probable overwintering of adult Hyalomma rufipes in Central Europe. Ticks Tick-Borne Dis. Munich: Elsevier Gmbh; 2021;12:101718.

10. Bah MT, Grosbois V, Stachurski F, Muñoz F, Duhayon M, Rakotoarivony I, et al. The Crimean-Congo haemorrhagic fever tick vector Hyalomma marginatum in the south of France: Modelling its distribution and determination of factors influencing its establishment in a newly invaded area. Transbound Emerg Dis. Wiley Online Library; 2022;69:e2351–65.

11. Krézsek C, Bally AW. The Transylvanian Basin (Romania) and its relation to the Carpathian fold and thrust belt: Insights in gravitational salt tectonics. Mar Pet Geol. Elsevier; 2006;23:405–42.

12. Karpelles L. Adalékok Magyarország atka-faunájához. Akadémia Kiadó; 1893.

13. Kotlán S. Adatok a hazai kullancs-fauna ismeretéhez [Data on the Hungarian tick fauna]. Állattani Közlöny. 1919;18:33–6.

14. Kotlán S. Adatok a hazai kullancs-fauna ismeretéhez [The Classification of the Ticks of Hungary]. Állattani Közlöny. 1921;20.

15. Földvári G, Szabó É, Tóth GE, Lanszki Z, Zana B, Varga Z, et al. Emergence of Hyalomma marginatum and Hyalomma rufipes adults revealed by citizen science tick monitoring in Hungary. Transbound Emerg Dis. Wiley Online Library; 2022;

16. Janisch M. Kullancsgazda madarak különféle betegségek közvetítői [Tick carrier birds spreading disease agents]. Aquila. 1960;67–68:191–4.

17. Csörgő T, Harnos A, Rózsa L, Karcza Z, Fehérvári P. Detailed description of the Ócsa Bird Ringing Station, Hungary. Ornis Hung. 2016;24:91–108.

18. Csörgő T, Karcza Z, Halmos G, Magyar G, Gyurácz J, Szép T, et al. Magyar madárvonulási atlasz [Atlas of bird migration in Hungary]. Budapest: Kossuth kiadó; 2009.

19. Folmer O, Black M, Hoeh W, Lutz R, Vrijenhoek R. DNA primers for amplification of mitochondrial cytochrome c oxidase subunit I from diverse metazoan invertebrates. Mol Mar Biol Biotechnol. 1994;3:294–9.

20. Black 4th WC, Piesman J. Phylogeny of hard-and soft-tick taxa (Acari: Ixodida) based on mitochondrial 16S rDNA sequences. Proc Natl Acad Sci. National Acad Sciences; 1994;91:10034–8.

21. Beati L, Keirans JE. Analysis of the systematic relationships among ticks of the genera Rhipicephalus and Boophilus (Acari: Ixodidae) based on mitochondrial 12S ribosomal DNA gene sequences and morphological characters. J Parasitol. 2001;87:32–48.

22. Bitencourth K, Voloch CM, Serra-Freire NM, Machado-Ferreira E, Amorim M, Gazêta GS. Analysis of Amblyomma sculptum haplotypes in an area endemic for Brazilian spotted fever. Med Vet Entomol. Wiley Online Library; 2016;30:342–50.

23. Burkman EJ. Genetic structure of Amblyomma cajennense (Acari: Ixodidae) populations based on mitochondrial gene sequences. 2009;

24. Hornok S, Karcza Z, Csörgő T. Birds as disseminators of ixodid ticks and tick-borne pathogens: note on the relevance to migratory routes. Ornis Hung. 2012;20:86–9.

25. Hornok S, Csörgő T, de la Fuente J, Gyuranecz M, Privigyei C, Meli ML, et al. Synanthropic birds associated with high prevalence of tick-borne rickettsiae and with the first detection of Rickettsia aeschlimannii in Hungary. Vector Borne Zoonotic Dis Larchmt N. 2013;13:77–83.

26. Hornok S, Flaisz B, Takács N, Kontschán J, Csörgő T, Csipak Á, et al. Bird ticks in Hungary reflect western, southern, eastern flyway connections and two genetic lineages of Ixodes frontalis and Haemaphysalis concinna. Parasit Vectors. 2016;9:101.

27. Wilhelmsson P, Jaenson TGT, Olsen B, Waldenstrom J, Lindgren P-E. Migratory birds as disseminators of ticks and the tick-borne pathogens Borrelia bacteria and tick-borne encephalitis (TBE) virus: a seasonal study at Ottenby Bird Observatory in South-eastern Sweden. Parasit Vectors. London: Bmc; 2020;13:607.

28. Heylen D, Fonville M, Docters van Leeuwen A, Stroo A, Duisterwinkel M, van Wieren S, et al. Pathogen communities of songbird-derived ticks in Europe’s low countries. Parasit Vectors. 2017;10:497.

29. Klaus C, Gethmann J, Hoffmann B, Ziegler U, Heller M, Beer M. Tick infestation in birds and prevalence of pathogens in ticks collected from different places in Germany. Parasitol Res. 2016;115:2729–40.

30. Toma L, Mancuso E, d’Alessio SG, Menegon M, Spina F, Pascucci I, et al. Tick species from Africa by migratory birds: a 3-year study in Italy. Exp Appl Acarol. Dordrecht: Springer; 2021;83:147–64.

31. Buczek AM, Buczek W, Buczek A, Bartosik K. The potential role of migratory birds in the rapid spread of ticks and tick-borne pathogens in the changing climatic and environmental conditions in Europe. Int J Environ Res Public Health. MDPI; 2020;17:2117.

32. Keve G, Sandor AD, Hornok S. Hard ticks (Acari: Ixodidae) associated with birds in Europe: Review of literature data. Front Vet Sci. Lausanne: Frontiers Media Sa; 2022;9:928756.

33. Janisch M. A hazai kullancsfauna feltérképezése [Geographical distribution of tick species in Hungary]. Állattani Közlöny. 1959;47:103–10.

34. Magano SR, Els DA, Chown SL. Feeding patterns of immature stages of Hyalomma truncatum and Hyalomma marginatum rufipes on different hosts. Exp Appl Acarol. Springer; 2000;24:301–13.

35. Hornok S, Cutajar B, Takács N, Galea N, Attard D, Coleiro C, et al. On the way between Africa and Europe: Molecular taxonomy of ticks collected from birds in Malta. Ticks Tick-Borne Dis. Elsevier; 2022;13:102001.

36. Márquez FJ, Caruz A. Phylogeography of Hyalomma (Euhyalomma) lusitanicum (Acarina, Parasitiformes, Ixodidae) in Andalusia based on mitochondrial cytochrome oxidase I gene. Exp Appl Acarol. Springer; 2021;85:49–61.

37. Hornok S. Allochronic seasonal peak activities of Dermacentor and Haemaphysalis spp. under continental climate in Hungary. Vet Parasitol. Elsevier; 2009;163:366–9.

38. Akimov IA, Nebogatkin IV. Distribution of Ticks from the Genus Dermacentor (Acari, Ixodidae) in Ukraine. Вестник Зоологии. Інститут зоології ім. ІІ Шмальгаузена НАН України; 2011;

39. Hubálek Z, Halouzka J, Juricova Z. Host-seeking activity of ixodid ticks in relation to weather variables. J Vector Ecol. Society for Vector Ecology; 2003;28:159–65.

40. Reynolds C, Kontschán J, Takács N, Solymosi N, Sándor AD, Keve G, et al. Shift in the seasonality of ixodid ticks after a warm winter in an urban habitat with notes on morphotypes of Ixodes ricinus and data in support of cryptic species within Ixodes frontalis. Exp Appl Acarol. Springer; 2022;88:127–38.

41. Pomerantsev BI. Ixodid Ticks (Ixodidae). American Institute of Biological Sciences; 1959.

42. Hoogstraal H. African Ixodoidea. VoI. I. Ticks of the Sudan (with special reference to Equatoria Province and with Preliminary Reviews of the Genera Boophilus, Margaropus, and Hyalomma). Afr Ixodoidea VoI Ticks Sudan Spec Ref Equat Prov Prelim Rev Genera Boophilus Margaropus Hyalomma [Internet]. 1956 [cited 2023 Jan 11]; Available from: https://www.cabdirect.org/cabdirect/abstract/19572901469

43. Wallmenius K, Barboutis C, Fransson T, Jaenson TGT, Lindgren P-E, Nystrom F, et al. Spotted fever Rickettsia species in Hyalomma and Ixodes ticks infesting migratory birds in the European Mediterranean area. Parasit Vectors. London: Bmc; 2014;7:318.

44. Kiros S, Awol N, Tsegaye Y, Hadush B. Hard Ticks of Camel in Southern Zone of Tigray, Northern Ethiopia. J Parasitol Vector Biol. 2014;6:151–5.

45. Kerario II, Muleya W, Chenyambuga S, Koski M, Hwang S-G, Simuunza M. Abundance and distribution of Ixodid tick species infesting cattle reared under traditional farming systems in Tanzania. Afr J Agric Res. Academic Journals; 2017;12:286–99.

