## Supplementary Figure 1 for "Ornithological and molecular evidence of a reproducing *Hyalomma rufipes* population under continental climate in Europe"

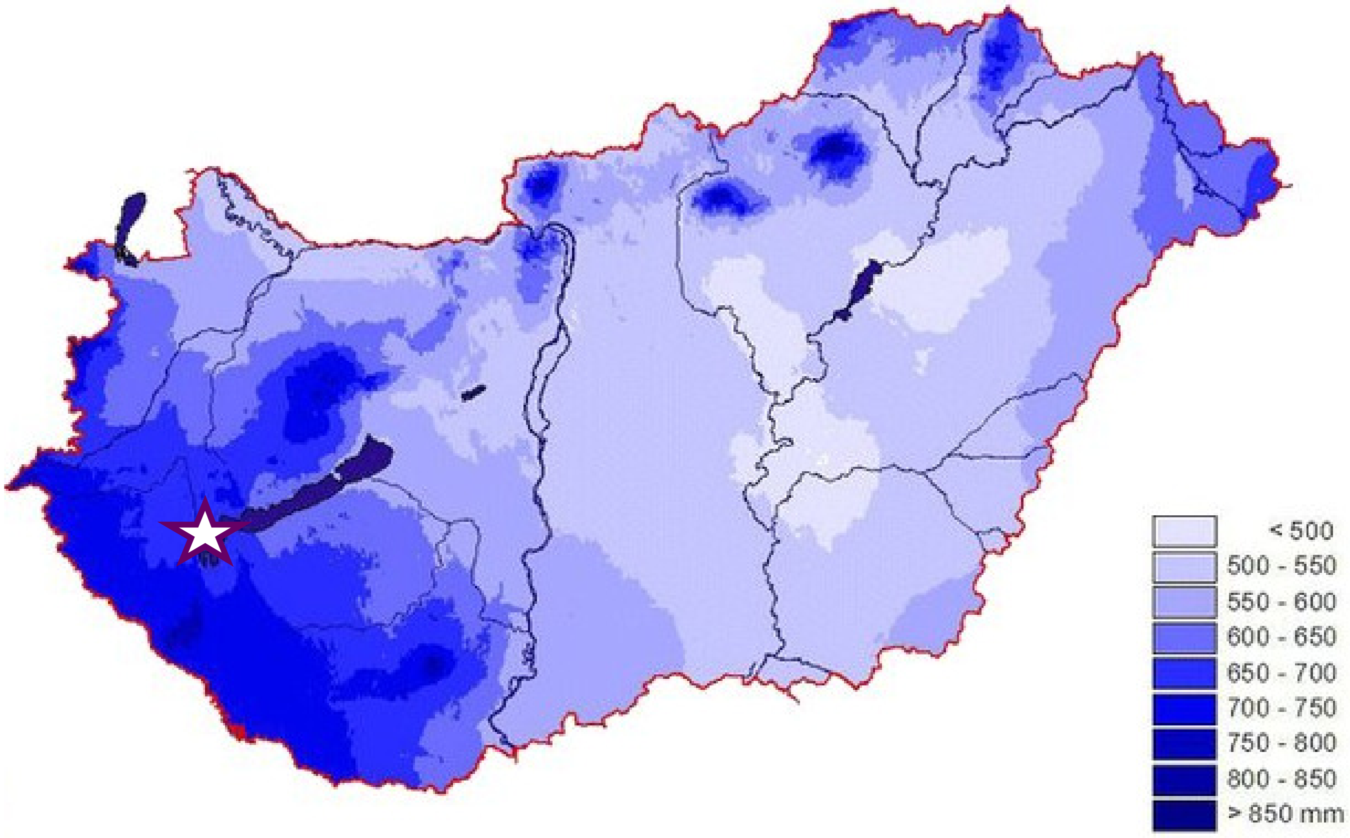


Average annual precipitation in Hungary (1971-2000).

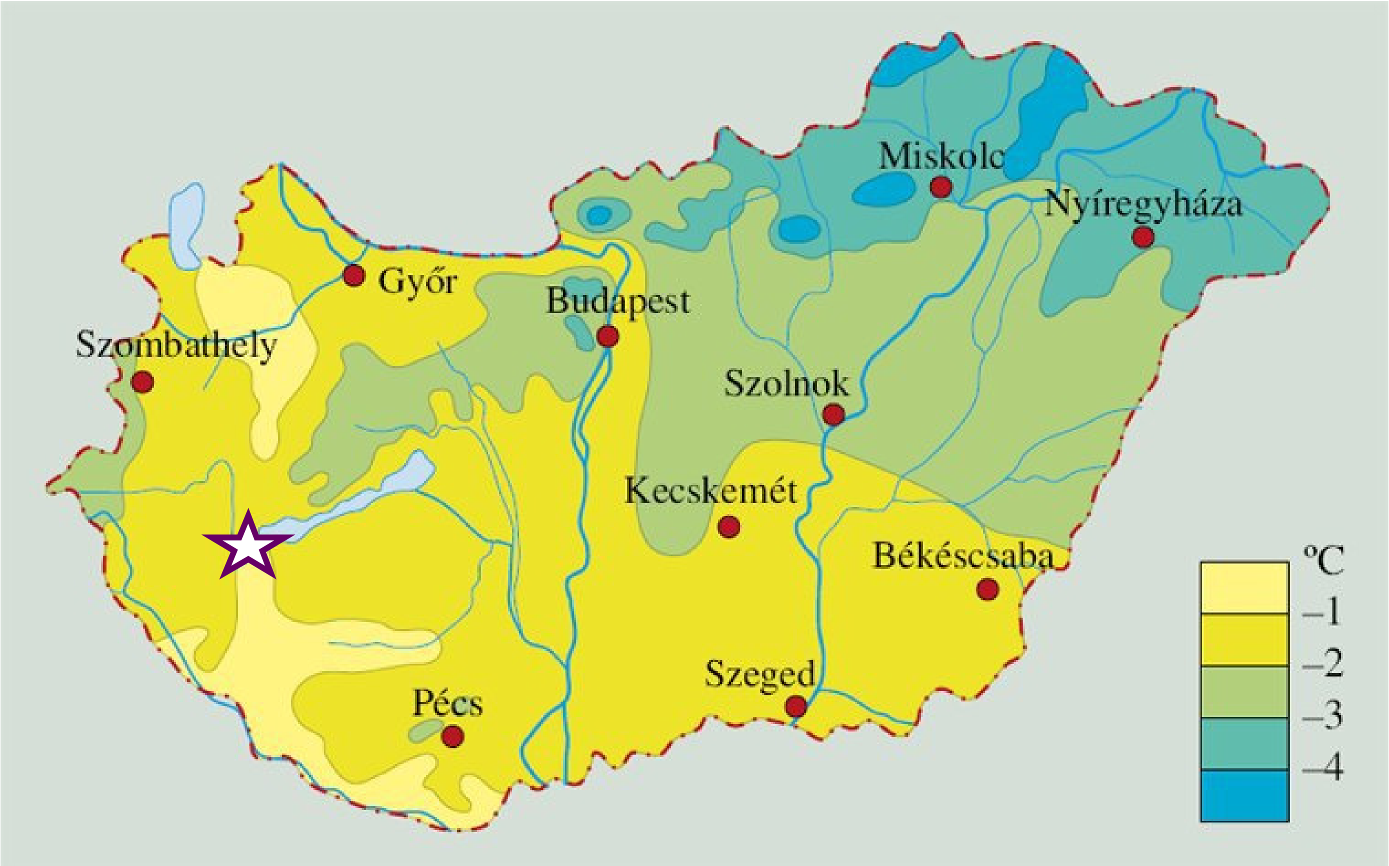


Average temperature in January in Hungary (data from the last two decades).
