## Supplementary Table 1 for "Ornithological and molecular evidence of a reproducing *Hyalomma rufipes* population under continental climate in Europe"

### Location: Dávod

#### Spring migration and summer nesting period (March-July):

| Tick species | Male | Female | Nymph | Larva | Total |
|---|---|---|---|---|---|
| *I. ricinus* | 0 | 0 | 15 | 10 | 25 |
| *H. concinna* | 0 | 0 | 30 | 5 | 35 |
| *Hy. marginatum* | 0 | 0 | 1 | 0 | 1 |
| *I. frontalis* | 0 | 0 | 0 | 0 | 0 |
| *I. lividus* | 0 | 6 | 0 | 0 | 6 |

#### Autumn migration period (August-November):

| Tick species | Male | Female | Nymph | Larva | Total |
|---|---|---|---|---|---|
| *I. ricinus* | 0 | 0 | 7 | 3 | 10 |
| *H. concinna* | 0 | 0 | 6 | 1 | 7 |
| *Hy. marginatum* | 0 | 0 | 0 | 0 | 0 |
| *I. frontalis* | 0 | 0 | 0 | 0 | 0 |

### Location: Lake Fehér

#### Spring migration and summer nesting period (March-July):

| Tick species | Male | Female | Nymph | Larva | Total |
|---|---|---|---|---|---|
| *I. ricinus* | 0 | 0 | 3 | 0 | 3 |
| *H. concinna* | 0 | 0 | 21 | 0 | 21 |
| *Hy. marginatum* | 0 | 0 | 0 | 0 | 0 |
| *I. frontalis* | 0 | 0 | 0 | 0 | 0 |

#### Autumn migration period (August-November):

| Tick species | Male | Female | Nymph | Larva | Total |
|---|---|---|---|---|---|
| *I. ricinus* | 0 | 0 | 0 | 1 | 1 |
| *H. concinna* | 0 | 0 | 11 | 3 | 14 |
| *Hy. marginatum* | 0 | 0 | 0 | 0 | 0 |
| *I. frontalis* | 0 | 0 | 0 | 0 | 0 |

### Location: Fenékpuszta, Lake Balaton

#### Spring migration and summer nesting period (March-July):

| Tick species | Male | Female | Nymph | Larva | Total |
|---|---|---|---|---|---|
| *I. ricinus* | 0 | 0 | 4 | 0 | 4 |
| *H. concinna* | 0 | 0 | 20 | 1 | 21 |
| *Hy. marginatum* | 0 | 0 | 9 | 1 | 10 |
| *I. frontalis* | 0 | 0 | 0 | 0 | 0 |

### Autumn migration period (August-November):

| Tick species | Male | Female | Nymph | Larva | Total |
|---|---|---|---|---|---|
| *I. ricinus* | 0 | 0 | 7 | 8 | 15 |
| *H. concinna* | 0 | 0 | 5 | 4 | 9 |
| *Hy. marginatum* | 0 | 0 | 0 | 0 | 0 |
| *I. frontalis* | 0 | 0 | 0 | 0 | 0 |

### Location: Izsák, Lake Kolon

### Spring migration and summer nesting period (March-July):

| Tick species | Male | Female | Nymph | Larva | Total |
|---|---|---|---|---|---|
| *I. ricinus* | 0 | 0 | 5 | 3 | 8 |
| *H. concinna* | 0 | 0 | 45 | 20 | 65 |
| *Hy. marginatum* | 0 | 0 | 0 | 0 | 0 |
| *I. frontalis* | 0 | 1 | 0 | 0 | 1 |

### Autumn migration period (August-November):

| Tick species | Male | Female | Nymph | Larva | Total |
|---|---|---|---|---|---|
| *I. ricinus* | 0 | 0 | 9 | 2 | 11 |
| *H. concinna* | 0 | 0 | 8 | 7 | 15 |
| *Hy. marginatum* | 0 | 0 | 0 | 0 | 0 |
| *I. frontalis* | 0 | 2 | 0 | 0 | 2 |

### Location: Ócsa

### Spring migration and summer nesting period (March-July):

| Tick species | Male | Female | Nymph | Larva | Total |
|---|---|---|---|---|---|
| *I. ricinus* | 0 | 0 | 121 | 2 | 123 |
| *H. concinna* | 0 | 0 | 62 | 55 | 117 |
| *Hy. marginatum* | 0 | 0 | 0 | 0 | 0 |
| *I. frontalis* | 0 | 0 | 1 | 0 | 1 |
| *D. reticulatus* | 0 | 1 | 0 | 0 | 1 |

### Autumn migration period (August-November):

| Tick species | Male | Female | Nymph | Larva | Total |
|---|---|---|---|---|---|
| *I. ricinus* | 0 | 0 | 50 | 34 | 84 |
| *H. concinna* | 0 | 0 | 5 | 3 | 8 |
| *Hy. marginatum* | 0 | 0 | 0 | 0 | 0 |
| *I. frontalis* | 0 | 1 | 1 | 3 | 5 |

### Location: Bódva Valley

#### Spring migration and summer nesting period (March-July):

| Tick species | Male | Female | Nymph | Larva | Total |
|---|---|---|---|---|---|
| *I. ricinus* | 0 | 0 | 0 | 0 | 0 |
| *H. concinna* | 0 | 0 | 0 | 0 | 0 |
| *Hy. marginatum* | 0 | 0 | 0 | 0 | 0 |
| *I. frontalis* | 0 | 0 | 0 | 0 | 0 |

#### Autumn migration period (August-November):

| Tick species | Male | Female | Nymph | Larva | Total |
|---|---|---|---|---|---|
| *I. ricinus* | 0 | 0 | 50 | 149 | 199 |
| *H. concinna* | 0 | 0 | 1 | 2 | 3 |
| *Hy. marginatum* | 0 | 0 | 0 | 0 | 0 |
| *I. frontalis* | 0 | 0 | 2 | 0 | 2 |

### Location: Tömörd

#### Spring migration and summer nesting period (March-July):

| Tick species | Male | Female | Nymph | Larva | Total |
|---|---|---|---|---|---|
| *I. ricinus* | 0 | 0 | 42 | 13 | 55 |
| *H. concinna* | 0 | 0 | 1 | 5 | 6 |
| *Hy. marginatum* | 0 | 0 | 1 | 0 | 1 |
| *I. frontalis* | 0 | 1 | 0 | 1 | 2 |

#### Autumn migration period (August-November):

| Tick species | Male | Female | Nymph | Larva | Total |
|---|---|---|---|---|---|
| *I. ricinus* | 0 | 0 | 44 | 16 | 60 |
| *H. concinna* | 0 | 0 | 1 | 0 | 1 |
| *Hy. marginatum* | 0 | 0 | 0 | 0 | 0 |
| *I. frontalis* | 0 | 4 | 0 | 1 | 5 |
