## Supplementary Table 2 for "Ornithological and molecular evidence of a reproducing *Hyalomma rufipes* population under continental climate in Europe"

**Supplementary Table 2.** Avian host species that were found tick-infested in this study, shown according to collection site and spring or autumn migration intervals (the former including nesting period). The number of tick-infested birds sampled at the same location is shown in parentheses after the abbreviation of bird species name. **Color code**: **red** - long distance migrant, **purple** - resident or short/mid-distance migrant, **blue** - resident.

### Spring migration and nesting (March to July):

| Tick species | Name of collection site | | | | | | |
|---|---|---|---|---|---|---|---|
| | **Tömörd** | **Ócsa** | **Bódva Valley** | **Fenékpuszta** | **Baja, Dávod** | **Lake Kolon** | **Lake Fehér** |
| *I. ricinus* | ERI RUB (8)<br>TUR MER (6)<br>TUR PHI (3)<br>PAR CAE (2)<br>PAR MAJ (5)<br>PAR PAL (1)<br>ACR RIS (1) | TUR MER (13)<br>PAR MAJ (4)<br>COC COC (1)<br>TUR PHI (7)<br>ERI RUB (6)<br>TRO TRO (4)<br>ACR SCH (2)<br>SYL ATR (4)<br>SYL COM (1)<br>ACR SCI (4)<br>PHY COL (2)<br>LUS MEG (3)<br>LOC LUS (4)<br>SIT EUR (1)<br>ACR MEL (1)<br>ACR RIS (4) | - | ACR SCI (3)<br>ACR ARU (1) | EMB SCH (2)<br>PAN BIA (1)<br>ACR SCI (6)<br>ACR ARU (1)<br>TUR MER (1)<br>TUR PHI (1)<br>SYL ATR (1)<br>LUS MEG (1)<br>ACR RIS (3) | ACR SCH (1)<br>ACR SCI (2)<br>ACR RIS (2)<br>PAR MAJ (1)<br>LUS MEG (1) | ACR SCI (1)<br>ACR RIS (1)<br>HYP ICT (1) |
| *H. concinna* | ERI RUB (3)<br>TUR PHI (1) | TUR MER (1)<br>TUR PHI (2)<br>ACR SCH (6)<br>ACR SCI (2)<br>LUS MEG (1)<br>LOC LUS (17)<br>ACR MEL (2)<br>ACR RIS (4) | - | ACR SCI (5)<br>ACR SCH (7)<br>EMB SCH (1)<br>LOC LUS (4) | ACR SCI (6)<br>ACR ARU (1)<br>ACR MEL (1)<br>TUR MER (1)<br>TUR PHI (1)<br>SYL ATR (1)<br>LOC LUS (11)<br>ACR SCH (3)<br>LOC NAE (1) | ACR SCH (25)<br>LOC LUS (17)<br>ACR SCI (4)<br>ACR RIS (4)<br>LOC NAE (2)<br>PAR CAE (1)<br>LOC FLU (1)<br>ACR ARU (1) | ACR SCI (3)<br>LOC LUS (6)<br>ACR SCH (3) |
| *Hyalomma* sp. | FIC HYP (1) | - | - | ACR SCH (3)<br>PAN BIA (2) | SYL COM (1) | - | - |
| *I. frontalis* | ERI RUB (2) | ACR SCI (1) | - | - | - | ACR SCI (1) | - |

### Autumn migration (August to November):

| Tick species | Name of collection site | | | | | | |
|---|---|---|---|---|---|---|---|
| | Tömörd | Ócsa | River Bódva | Fenékpuszta | Baja, Dávod | Lake Kolon | Lake Fehér |
| *I. ricinus* | ERI RUB (7)<br>TUR MER (17)<br>TUR PHI (1)<br>PAR MAJ (1)<br>ANT TRI (1)<br>LUS MEG (1)<br>SYL ATR (4)<br>PHY TRO (1)<br>SYL COM (2)<br>PAS MON (2)<br>PHO OCH (1)<br>CAR CHL (1)<br>TUR ILI (1)<br>TRO TRO (2) | TUR MER (7)<br>PAR MAJ (1)<br>TUR PHI (2)<br>ERI RUB (16)<br>ACR SCH (1)<br>SYL ATR (8)<br>SYL COM (5)<br>ACR SCI (1)<br>LUS MEG (5)<br>ACR RIS (3)<br>SYL BOR (2)<br>LUS LUS (7)<br>CER BRA (1)<br>FRI COE (1) | ERI RUB (62)<br>SYL ATR (8)<br>SYL COM (6)<br>LUS MEG (1)<br>LUS LUS (1)<br>TUR MER (11)<br>FRI COE (1)<br>TUR PHI (2)<br>PHY TRO (1) | ACR SCI (2)<br>ACR SCH (2)<br>SYL ATR (2)<br>SYL COM (2)<br>HIR RUS (1)<br>LUS LUS (1)<br>PHY TRO (1)<br>LUS SVE (1)<br>ERI RUB (2) | SYL COM (1)<br>ACR SCI (1)<br>TUR MER (2)<br>SYL ATR (1)<br>ERI RUB (1)<br>TRO TRO (2)<br>ACR RIS (2) | LOC LUS (1)<br>LUS LUS (2)<br>ERI RUB (3)<br>TUR MER (1) | ACR RIS (1) |
| *H. concinna* | TUR MER (1) | TUR PHI (1)<br>ACR SCI (3)<br>LUS MEG (1)<br>LOC LUS (1) | ERI RUB (1)<br>SYL ATR (2) | ACR SCI (2)<br>ACR SCH (4)<br>ACR ARU (1)<br>LOC FLU (1) | ACR SCI (1)<br>TUR MER (1)<br>LOC LUS (3)<br>ACR SCH (1)<br>ACR RIS (1) | ACR ARU (1)<br>LOC LUS (5)<br>ACR RIS (1)<br>LOC NAE (1) | ACR SCI (1)<br>LOC LUS (1)<br>ACR ARU (2) |
| *Hyalomma* sp. | - | - | - | - | - | - | - |
| *I. frontalis* | ERI RUB (1)<br>TUR MER (2)<br>PAS MON (1) | TUR PHI (1)<br>ERI RUB (1)<br>PAS MON (1) | ERI RUB (1)<br>TUR MER (1) | - | - | ACR SCI (1)<br>TUR MER (1) | - |

**Abbreviations:**

ACR RIS = *Acrocephalus palustris*, ACR SCH = *A. schoenobaenus*, ACR SCI = *A. scirpaceus*, LOC LUS = *Locustella luscinioides*, LOC NAE = *L. naevia*, PHY COL = *Phylloscopus collybita*, SYL ATR = *Sylvia atricapilla*, CAR CHL = *Carduelis chloris*, COC COC = *Coccothraustes coccothraustes*, EMB CIT = *Emberiza citrinella*, EMB SCH = *E. schoeniclus*, PAR MAJ = *Parus major*, LUS LUS = *Luscinia luscinia*, LUS MEG = *L. megarhynchos*, SYL COM = *S. communis*, TUR ILI = *Turdus iliacus*, ERI RUB = *Erithacus rubecula*, PRU MOD = *Prunella modularis*, TRO TRO = *Troglodytes troglodytes*, TUR MER = *T. merula*, TUR PHI = *T. philomelos*
